# BMP signaling to pharyngeal muscle in the *C. elegans* response to a bacterial pathogen regulates anti-microbial peptide expression and pharyngeal pumping

**DOI:** 10.1101/2023.03.06.531324

**Authors:** Emma Jo Ciccarelli, Moshe Bendelstein, Katerina K. Yamamoto, Hannah Reich, Cathy Savage-Dunn

## Abstract

Host response to pathogens recruits multiple tissues in part through conserved cell signaling pathways. In *C. elegans*, the bone morphogenetic protein (BMP) like DBL-1 signaling pathway has a role in the response to infection in addition to other roles in development and post-developmental functions. In the regulation of body size, the DBL-1 pathway acts through cell autonomous signal activation in the epidermis (hypodermis). We have now elucidated the tissues that respond to DBL-1 signaling upon exposure to two bacterial pathogens. The receptors and Smad signal transducers for DBL-1 are expressed in pharyngeal muscle, intestine, and epidermis. We demonstrate that expression of receptor-regulated Smad (R-Smad) gene *sma-3* in the pharynx is sufficient to improve the impaired survival phenotype of *sma-3* mutants and that expression of *sma-3* in the intestine has no effect when exposing worms to bacterial infection of the intestine. We also show that two antimicrobial peptide genes – *abf-2* and *cnc-2* – are regulated by DBL-1 signaling through R-Smad SMA-3 activity in the pharynx. Finally, we show that pharyngeal pumping activity is reduced in *sma-3* mutants and that other pharynx-defective mutants also have reduced survival on a bacterial pathogen. Our results identify the pharynx as a tissue that responds to BMP signaling to coordinate a systemic response to bacterial pathogens.

**Significance Statement:**

- Innate immunity is the first line of defense against pathogens. Conserved cell signaling pathways are known to be involved in host-pathogen response, but how they coordinate a systemic response is less well understood.
- In the nematode *C. elegans,* bone morphogenetic protein (BMP) signaling is required for survival on pathogenic bacteria. Using transgenic strains, the authors identify a major role for a specific organ, the pharynx, in BMP-dependent survival.
- These findings demonstrate that an organ can serve as a pathogen sensor to trigger multiple modes of response to bacterial pathogens, include a barrier response and regulation of anti-microbial peptide expression.

## Introduction

Animals encounter a diverse range of microorganisms in their environments, many of which can pose a risk for infection. This exposure has resulted in the development of immune networks to protect the host from pathogenic threats. Innate immunity provides an immediate response to infection and functions through highly conserved mechanisms. In humans, innate immunity involves physical barriers such as the skin, recruitment of phagocytic cells, and the release of antimicrobial peptides (AMPs). Expression of these AMPs is regulated by a number of signaling pathways and these mechanisms are conserved across metazoan species (Medzhitov and Janeway, 1998; Hoffman et al., 1999; Kim et al., 2002; Schulenburg et al., 2004; Millet and Ewbank, 2004; Irazoqui et al., 2008; Partridge et al., 2010; Pukkila-Worley and Ausubel, 2012; Kim and Ewbank, 2018).

The soil-dwelling nematode *Caenorhabditis elegans* naturally encounters many pathogens, thus necessitating a functioning immune system to provide adequate protection. Pathogen response can be divided into behavioral avoidance responses, innate immunity (including barrier functions and AMP expression), and immune tolerance (mitigating the effect of infection rather than reducing infection) (Yamamoto and Savage-Dunn, 2023). The initial point of contact for pathogens can be the cuticle, intestine, uterus, or rectum (Kim and Ewbank, 2018). In addition, defects in pharyngeal function can cause increased susceptibility to ingested microbes (Labrousse et al., 2000; Kim et al., 2002). Although the animals lack an antibody-based acquired immunity, *C. elegans* has served as a useful organism in which to study many of the conserved signaling pathways that are involved in innate immunity and how they regulate an antimicrobial response (Kim et al., 2002; Schulenburg et al., 2004; Millet and Ewbank, 2004; Irazoqui et al., 2008; Partridge et al., 2010; Pukkila-Worley and Ausubel, 2012; Kim and Ewbank, 2018).

One pathway involved in the *C. elegans* immune response is the BMP-like DBL-1 signaling pathway (Mallo et al., 2002). BMPs are members of the large conserved Transforming Growth Factor beta (TGF-β) family of secreted peptide growth factors (Massagué, 1998). Members of the TGF-β family play critical roles in adaptive and innate immunity (Chen and ten Dijke, 2016). In *C. elegans,* the DBL-1 pathway (Fig 1A) is initiated through the neuronally expressed ligand DBL-1 (Suzuki et al., 1999; Morita et al., 2002). The receptors DAF-4 and SMA-6 along with the signaling mediators SMA-2, SMA-3, and SMA-4 are endogenously expressed in the epidermis (also known as hypodermis in *C. elegans*), the intestine, and the pharynx (Savage et al., 1996; Suzuki et al., 1999; Inoue and Thomas, 2000; Savage-Dunn et al., 2000; Yoshida et al., 2001; Mallo et al., 2002; Wang et al., 2002). DBL-1 signaling plays a significant role in body size regulation, male tail development, mesodermal patterning, and lipid accumulation (Attisano and Wrana, 1996; Savage et al., 1996; Heldin et al., 1997; Derynck et al., 1998; Padgett et al., 1998; Whitman, 1998; Suzuki et al., 1999; Inoue and Thomas, 2000; Savage-Dunn et al., 2000; Yoshida et al., 2001; Wang et al., 2002; Gumienny and Savage-Dunn, 2018; Clark et al., 2018) in addition to its role in the *C. elegans* immune response (Mallo et al., 2002). This pathway has been well studied in the *C. elegans* response to the Gram-negative bacterium *Serratia marcescens,* where it has been demonstrated that mutation in *dbl-1* results in increased susceptibility to infection (Mallo et al., 2002). Our interest in this study was to determine the way in which DBL-1 signaling plays a broader role in the immune response against bacterial pathogens and the mechanisms through which this specific function is carried out.

**Figure 1.**
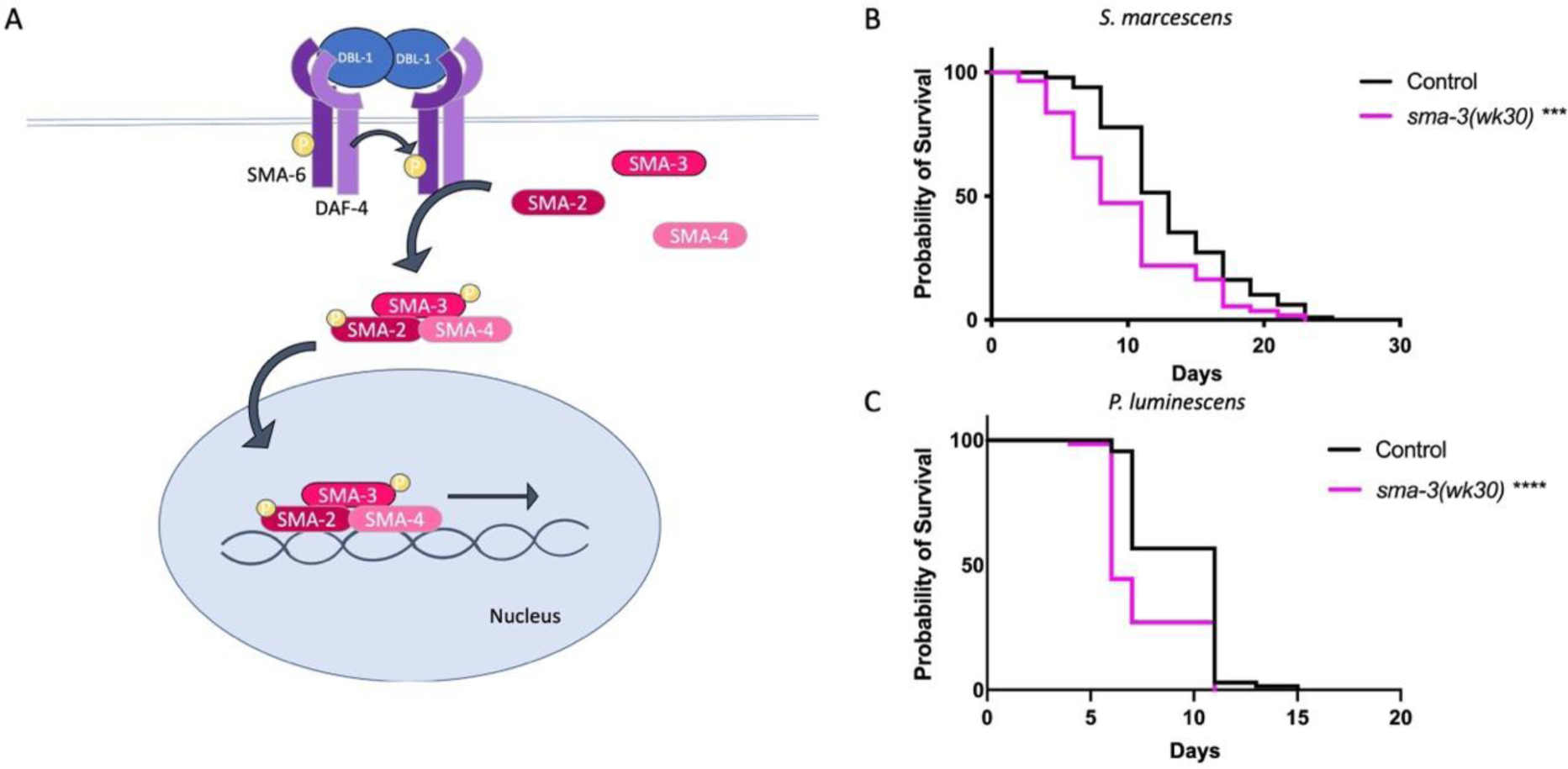
DBL-1 Pathway R-Smad SMA-3 is required for survival on bacterial pathogen. **A.** BMP signaling in *C. elegans* through the DBL-1 pathway. **B.** Survival of *sma-3(wk30)* on *S. marcescens.* n values: Control (99), *sma-3* (55). **C.** Survival of *sma-3(wk30)* on *P. luminescens*. n values: Control (67), *sma-3* (63). Survival analyses were repeated and results were consistent across trials. Statistical analysis was done using Log-rank (Mantel-Cox) Test. *** P ≤ 0.001; **** P < 0.0001.

Here we show that DBL-1 signaling through the receptor-regulated R-Smad SMA-3 is required for protection against bacterial pathogen in *C. elegans*. We demonstrate that expression of *sma-3* in pharyngeal muscle is sufficient to restore survival of *sma-3* deficient animals. DBL-1 signaling regulates expression of antimicrobial peptides ABF-2 (antibacterial factor related) (Kato et al., 2002) and CNC-2 (Caenacin) (Zugasti and Ewbank, 2009) and this regulation is through R-Smad SMA-3. We further show that pharyngeal expression of SMA-3 regulates ABF-2 and CNC-2 expression following pathogen exposure. Finally, we demonstrate that *sma-3* mutants have a reduced pharyngeal pumping rate that could increase sensitivity to pathogens and that expression of *sma-3* in pharyngeal muscle restores the pumping rate to control levels.

## Results

### DBL-1 Pathway R-Smad SMA-3 is required for survival against bacterial pathogen

Canonical TGF-β signaling employs a Co-Smad that forms a heterotrimer with R-Smads (Massagué, 1998). In response to fungal infection with *D. coniospora*, DBL-1/BMP signaling provides host protection through a non-canonical signaling cascade that does not include a Co-Smad (Zugasti and Ewbank, 2009). DBL-1 signaling activity is also required for *C. elegans* survival against infection by the Gram-negative bacterium *S. marcescens* (Mallo et al., 2002); which signaling components are required for this response was not determined. We therefore analyzed the survival phenotype of R-Smad mutant *sma-3* on *S. marcescens*. Animals were transferred from *E. coli* food to pathogenic bacteria at the fourth larval (L4) stage and survival on pathogen was monitored through adulthood until death. When we compare the survival of *sma-3* mutant animals, we see that they display a significantly reduced survival compared to control animals when exposed to *S. marcescens* infection, thus indicating that SMA-3 is involved in the response to *S. marcescens* infection (Fig 1B). We similarly observed that *sma-3* mutants have reduced survival when infected with the Gram-negative bacterium *P. luminescens* (Fig 1C), a pathogenic bacterium that is more virulent than *S. marcescens* against *C*. *elegans.* We have previously shown that *sma-3* mutants do not have a shortened lifespan on non-pathogenic bacteria (Clark et al., 2021). We conclude that SMA-3 plays a significant role in *C. elegans* survival against different bacterial pathogens.

### Expression of *sma-3(+)* in pharyngeal muscle rescues survival of *sma-3* mutants

DBL-1 pathway Smads – SMA-2, SMA-3, and SMA-4 – are endogenously expressed in the epidermis, the intestine, and the pharynx (Savage et al., 1996; Suzuki et al., 1999; Inoue and Thomas, 2000; Savage-Dunn et al., 2000; Yoshida et al., 2001; Mallo et al., 2002; Wang et al., 2002). We have previously used tissue-specific rescue of *sma-3* mutants to address its focus of action for different biological functions. For body size regulation, expression of *sma-3* in the epidermis alone is sufficient to rescue the body size phenotype in a *sma-3* mutant background (Wang et al., 2002).

We now sought to address in which tissue(s) SMA-3 activity is required to provide protection against bacterial infection. We used transgenic *C. elegans* strains in a *sma-3* mutant background to conduct these experiments. These strains have *sma-3* expressed under the control of tissue-specific promoters (control using endogenous *sma-3p* promoter; intestinal expression using *vha-6p*; pharyngeal muscle expression using *myo-2p*; epidermal expression using *vha-7p*), allowing for expression of *sma-3* in a tissue-specific manner. We hypothesized that intestinal SMA-3 would rescue survival, as the intestine is the main tissue in which an immune response to pathogenic bacteria has been described. Upon exposure to *S. marcescens* we found that expression of *sma-3* in pharyngeal muscle but not in intestine resulted in rescue of the *sma-3* mutant reduced survival phenotype. (Fig 2A). We repeated this experiment using *P. luminescens* and similarly found that pharyngeal expression of *sma-3* alone was sufficient to rescue the *sma-3* mutant survival phenotype (Fig 2B). Interestingly, on this pathogen, intestinal expression of *sma-3* resulted in a worse survival outcome compared to *sma-3* mutants. In initial trials it appeared that epidermal expression also rescued survival, but further examination revealed that the strain being employed had expression in additional tissues as well. We therefore repeated the survival experiments with a new, validated epidermal-specific strain (Supplemental Figure S1). In this strain, epidermal expression of *sma-3* failed to rescue (Supplemental Figure S2). From these results we conclude that the pharynx plays a significant role for DBL-1 pathway activity in response to bacterial infection in *C. elegans*.

**Figure 2.**
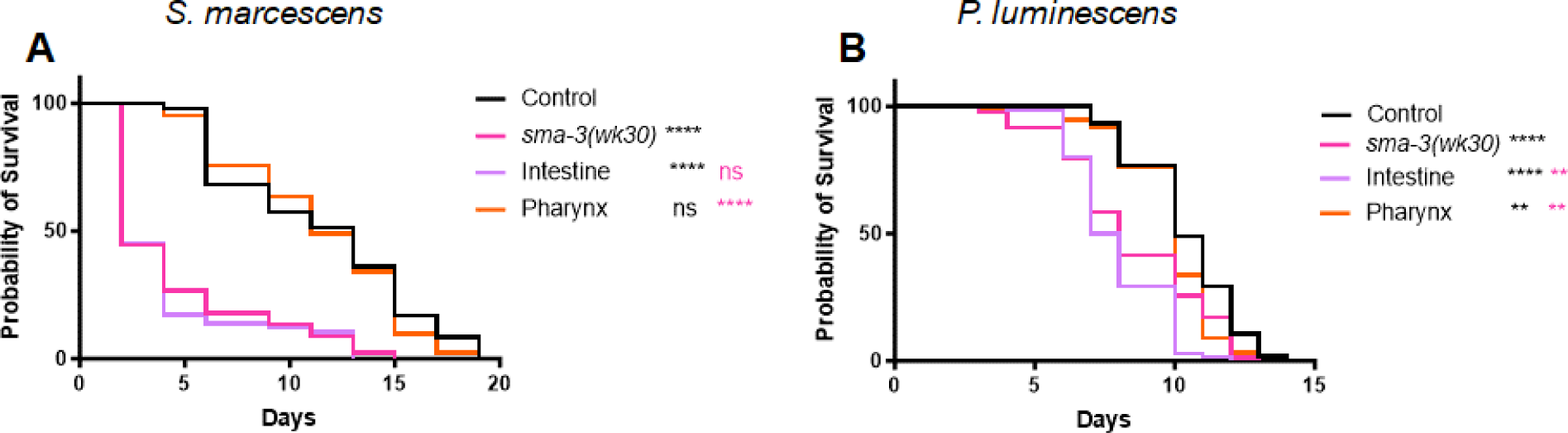
Expression of *sma-3(+)* in pharyngeal muscle rescues survival of *sma-3* mutants. **A.** Survival analysis of tissue-specific *sma-3* expressing strains on *S. marcescens*. Tissue-specific strains have *sma-3* expressed under the regulation of tissue-specific promotors in a *sma-3(wk30)* mutant background: *vha-6p* for intestinal expression and *myo-2p* for pharyngeal expression. n values: Control (47), *sma-3* (45) Intestine (58), Pharynx (41). **B.** Survival of tissue-specific *sma-3* expressing strains on *P. luminescens.* n values: Control (102), *sma-3* (82), Intestine (134), Pharynx (131). Survival analyses were repeated and results were consistent across trials. Statistical analysis was done using Log-rank (Mantel-Cox) Test. ns P > 0.05; * P ≤ 0.05; ** P ≤ 0.01; *** P ≤ 0.001; **** P < 0.0001. Black asterisks show significance relative to control; magenta shows significance relative to *sma-3* mutant.

### DBL-1 regulates expression of AMPs *abf-2* and *cnc-2* through SMA-3

We next tested whether the protective nature of DBL-1 signaling against bacterial infection is due to induction of AMP expression. Because a transcriptional response to *P. luminescens* is detected after 24 hours of exposure, while the transcriptional response to *Serratia* takes longer, we performed transcriptional assays in response to *P. luminescens.* We tested a panel of 12 immune response genes via qRT-PCR to determine whether DBL-1 signaling regulates their expression when animals are exposed to *P. luminescens* (Supplemental Figure S3). L4 animals were transferred from control plates to pathogen and collected as Day 1 adults following 24-hour exposure to pathogen. From this analysis we found two genes that demonstrated the expression pattern we expected if their regulation was mediated through DBL-1. Both *abf-2* and *cnc-2* were significantly upregulated in control animals upon 24-hour exposure to bacterial pathogen. Neither of these genes had a significant increase in expression in *dbl-1* mutant animals under the same infection conditions (Fig 3A).

**Figure 3.**
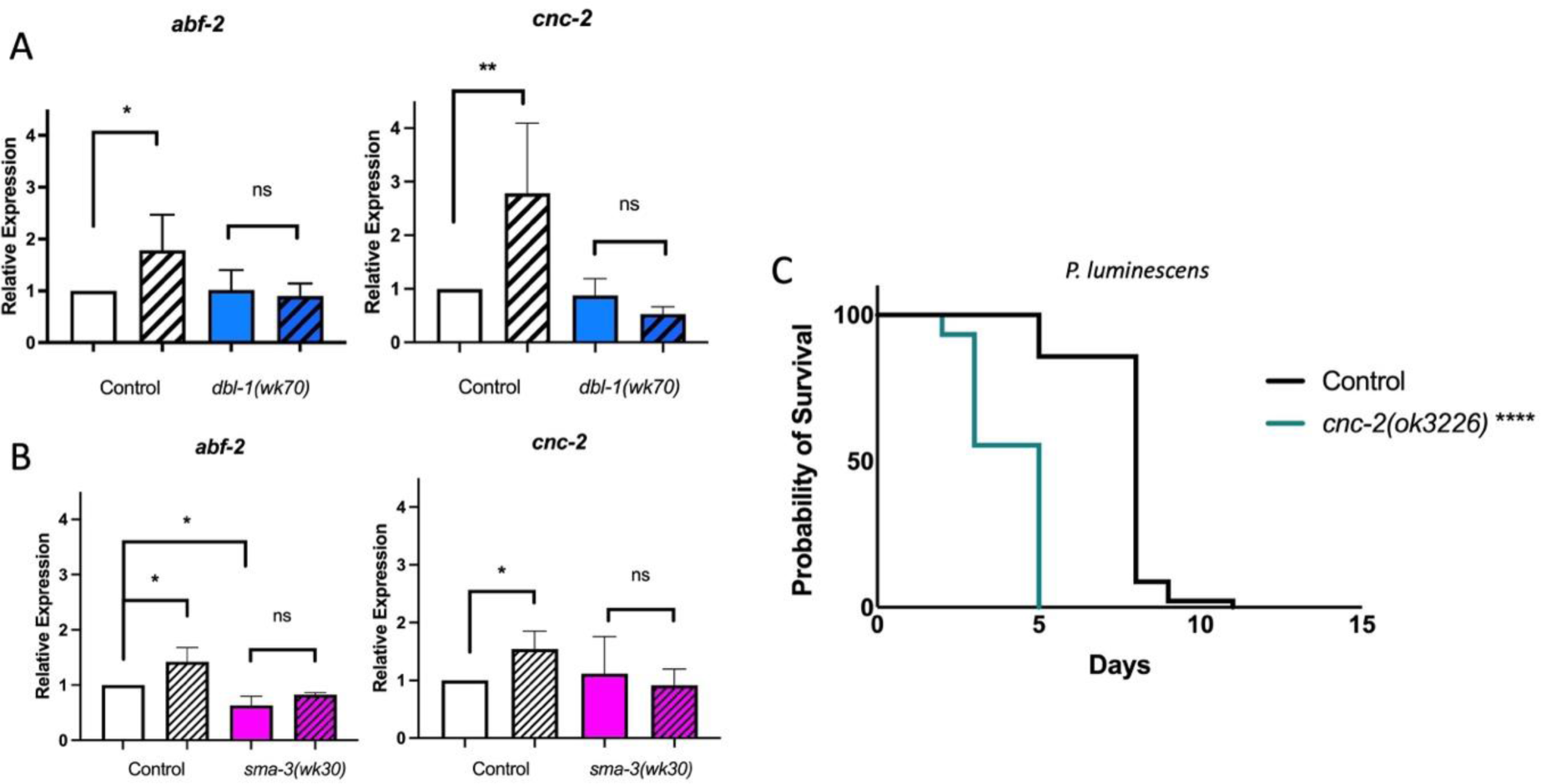
DBL-1 regulates expression of AMPs *abf-2* and *cnc-2* through SMA-3. **A**. qRT-PCR results for *abf-2* and *cnc-2* expression in *dbl-1(wk70)* animals on control bacteria and *P. luminescens*. **B.** qRT-PCR results for *abf-2* and *cnc-2* in *sma-3(wk30)* animals on control bacteria and *P. luminescens*. **C.** Survival analysis of *cnc-2(ok3226)* animals on *P. luminescens* pathogenic bacteria. n values: Control (91), *cnc-2* (74). Survival analysis was repeated and results were consistent across trials. Data for qRT-PCR represents repeated analyses from two biological replicates. Statistical analysis for qRT-PCR was done using One-way ANOVA with Šídák’s multiple comparison test using GraphPad Prism 9. Error bars represent standard deviation. Statistical analysis for survival was done using Log-rank (Mantel-Cox) Test. ns P > 0.05; * P ≤ 0.05; ** P ≤ 0.01; **** P < 0.0001.

ABF-2 is an antibacterial factor related protein that has been shown to be expressed in the pharynx of *C. elegans* and is involved in the response against bacterial infection (Kato et al., 2002). *cnc-2* encodes a Caenacin protein previously shown to be involved in the *C. elegans* immune response to fungal infection, where it is regulated in epidermal tissue (Zugasti and Ewbank, 2009). Our results suggest that both *abf-2* and *cnc-2* are regulated by DBL-1 signaling during the response to a bacterial pathogen.

We then used qRT-PCR to test whether *sma-3* mutants demonstrate any change in *abf-2* levels when exposed to bacterial pathogen. Following 24-hour exposure to *P. luminescens* bacteria, *sma-3* mutants demonstrated no increase in *abf-2* levels as compared to control conditions. Additionally, *sma-3* mutants expressed overall lower levels of *abf-2* than control animals even in control conditions (Fig 3B). This observation indicates that SMA-3 plays a role in the regulation of antimicrobial peptide ABF-2. When we use qRT-PCR to assess the expression pattern of *cnc-2* in *sma-3* mutants, we find a similar pattern for *cnc-2* expression. Control animals have a significant increase in *cnc-2* expression under infection conditions while *sma-3* mutants demonstrate no significant change in *cnc-2* expression when exposed to bacterial pathogen (Fig 3B). From these data we conclude that DBL-1 signaling does regulate expression of known immune response genes and that loss of the signaling effector SMA-3 prevents induction of both *abf-2* and *cnc-2* expression.

As previously noted, earlier work showed that *cnc-2* is involved in the *C. elegans* response to fungal infection (Zugasti and Ewbank, 2009). *cnc-2* was shown to be upregulated in response to *Serratia* as well (Mallo et al., 2002), but that study did not test its function in survival. We therefore followed up these experiments with a survival analysis of a *cnc-2* mutant on *P. luminescens* to determine whether *cnc-2* plays a role in bacterial infection as well as fungal infection. We used *cnc-2(ok3226)* mutants, a likely null allele with a 509 bp deletion of the *cnc-2* locus. *cnc-2* mutant animals had a significantly reduced survival rate when exposed to *P. luminescens* as compared to control animals (Fig 3C). We therefore conclude that CNC-2 plays a role in bacterial infection, in addition to its known role against fungal pathogen (Zugasti and Ewbank, 2009).

### DBL-1 pathway mediated regulation of AMP expression requires pharyngeal SMA-3 activity

Bacterial infection such as that caused by *S. marcescens* or *P. luminescens* results in colonization in the intestine of the worm (Tan et al., 1999; Garsin et al., 2001; Mallo et al., 2002; Kurz and Ewbank, 2003; Sato et al., 2014). As our survival analyses demonstrated that intestine-specific *sma-3* expression was not sufficient to ameliorate the *sma-3* reduced survival phenotype, we next tested whether DBL-1 mediated regulation of antimicrobial factors depends on pharyngeal SMA-3. We performed qRT-PCR analysis to establish the expression patterns for *abf-2* and *cnc-2* after a 24-hour exposure to *P. luminescens* bacteria. Consistent with our survival analysis results, which indicated significance for pharyngeal *sma-3* expression in *C. elegans* survival, *sma-3* expression in the pharynx was sufficient to cause an induction in *abf-2* and *cnc-2* transcript levels following *P. luminescens* exposure (Fig 4AB).

**Figure 4.**
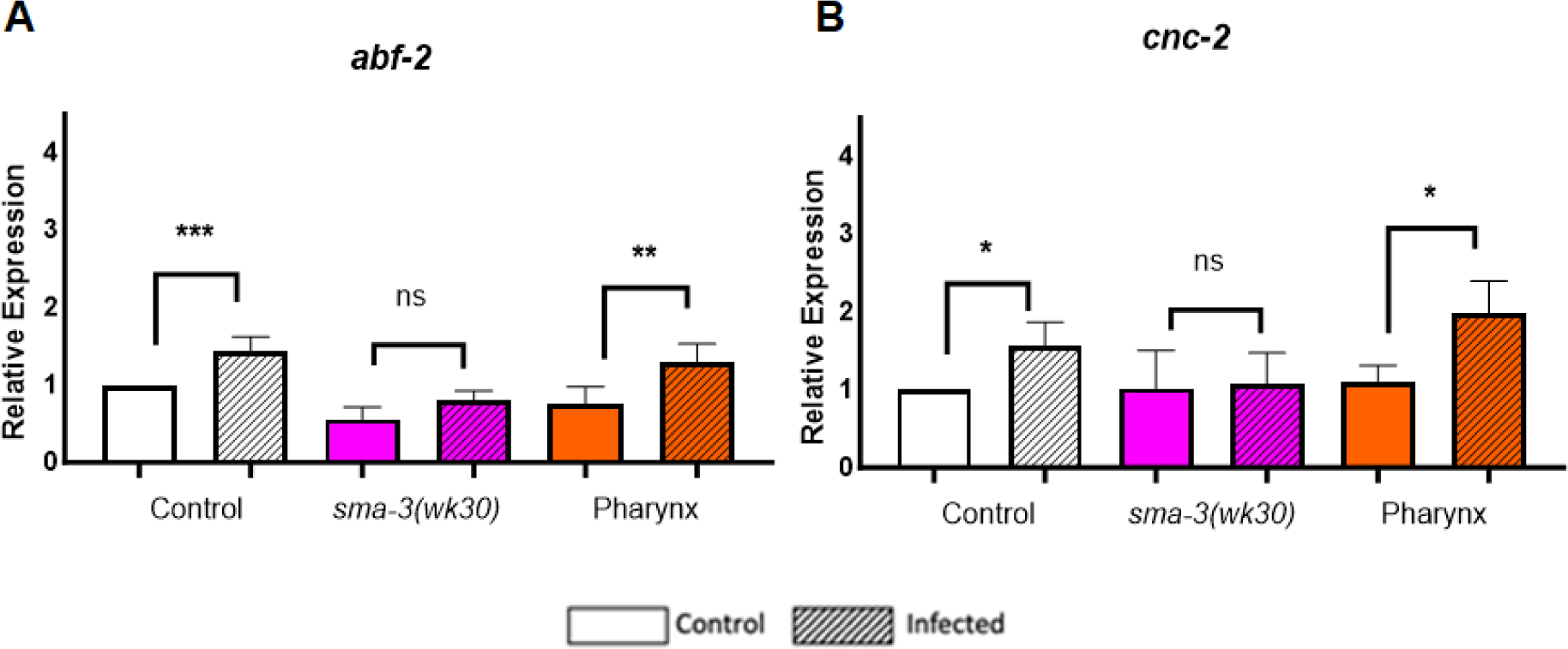
Pharyngeal expression of SMA-3 mediates regulation of AMP expression. **A.** qRT-PCR results for *abf-2* in transgenic *sma-3* expressing animals on control bacteria and *P. luminescens*. **B.** qRT-PCR results for *cnc-2* in transgenic *sma-3* expressing animals on control bacteria and *P. luminescens*. Data for qRT-PCR represents repeated analyses from two biological replicates. Statistical analysis for qRT-PCR was done using One-way ANOVA with Šídák’s multiple comparison test, using GraphPad Prism 9. Error bars represent standard deviation. ns P > 0.05; * P ≤ 0.05; ** P ≤ 0.01; *** P < 0.001.

### Expression of *sma-3* in pharyngeal muscle restores pumping rate in *sma-3* mutants

We next investigated whether *sma-3* expression in the pharynx resulted in any effect on the mechanical function of the pharynx, by assaying the pharyngeal pumping rate. To match the conditions of our survival and gene expression studies, we transferred L4 animals from control plates to plates seeded with pathogenic *P. luminescens* bacteria and measured pumping rates in Day 1 adults after 24 hours of exposure to pathogen. We found that *sma-3* mutants had a reduction in pumping rate as compared to control animals on pathogen. The pumping rate of *sma-3* mutants was restored when *sma-3* was expressed solely in the pharynx (Fig 5A), indicating potentially cell autonomous regulation of pumping rate by SMA-3. We also determined the pharyngeal pumping rate of *sma-3* mutants on nonpathogenic *E. coli*, and found that it was similarly reduced compared with wild type (Fig 5B). This result indicates that the reduced pumping rate is an underlying property of *sma-3* mutants, not a pathogen-induced alteration. Reduced pumping rate could potentially have two opposing effects; either lowering intake of bacteria, which would be expected to increase survival, or allowing intake of more intact, live bacteria, which would be more consistent with the observed reduced survival. To distinguish between these alternatives, we fed animals fluorescently tagged OP50-GFP (nonpathogenic *E. coli*) and imaged them after 24 hours of exposure (Fig 5 C-F). As expected, wild-type N2 animals display no fluorescent bacteria in their intestinal lumens after 24-hour exposure to OP50-GFP (Fig 5C). A small amount of fluorescent bacteria can be seen in the pharynx, where it is likely destroyed in the grinder. In contrast, a pharyngeal defective mutant, *phm-2*, displays a significant accumulation of bacteria in their intestinal lumens after 24-hour exposure to OP50-GFP (Fig 5D), consistent with a failure to destroy bacteria in the pharynx. *dbl-1* mutants have slightly more bacterial accumulation than wild type after 24-hour exposure to OP50-GFP (Fig 5E). However, levels remain low compared to *phm-2* mutants. This finding is consistent with the normal pumping rate of *dbl-1* mutants on *E. coli* and *P. luminescens* (Ciccarelli et al., 2023). The slight accumulation of live bacteria is likely due to the innate immune defect of this mutant. Finally, *sma-3* mutants display very high bacterial accumulation after 24-hour exposure to OP50-GFP (Fig 5F), similar to that in *phm-2* mutants, demonstrating the consequences of the grinder defect in *sma-3* animals.

**Figure 5.**
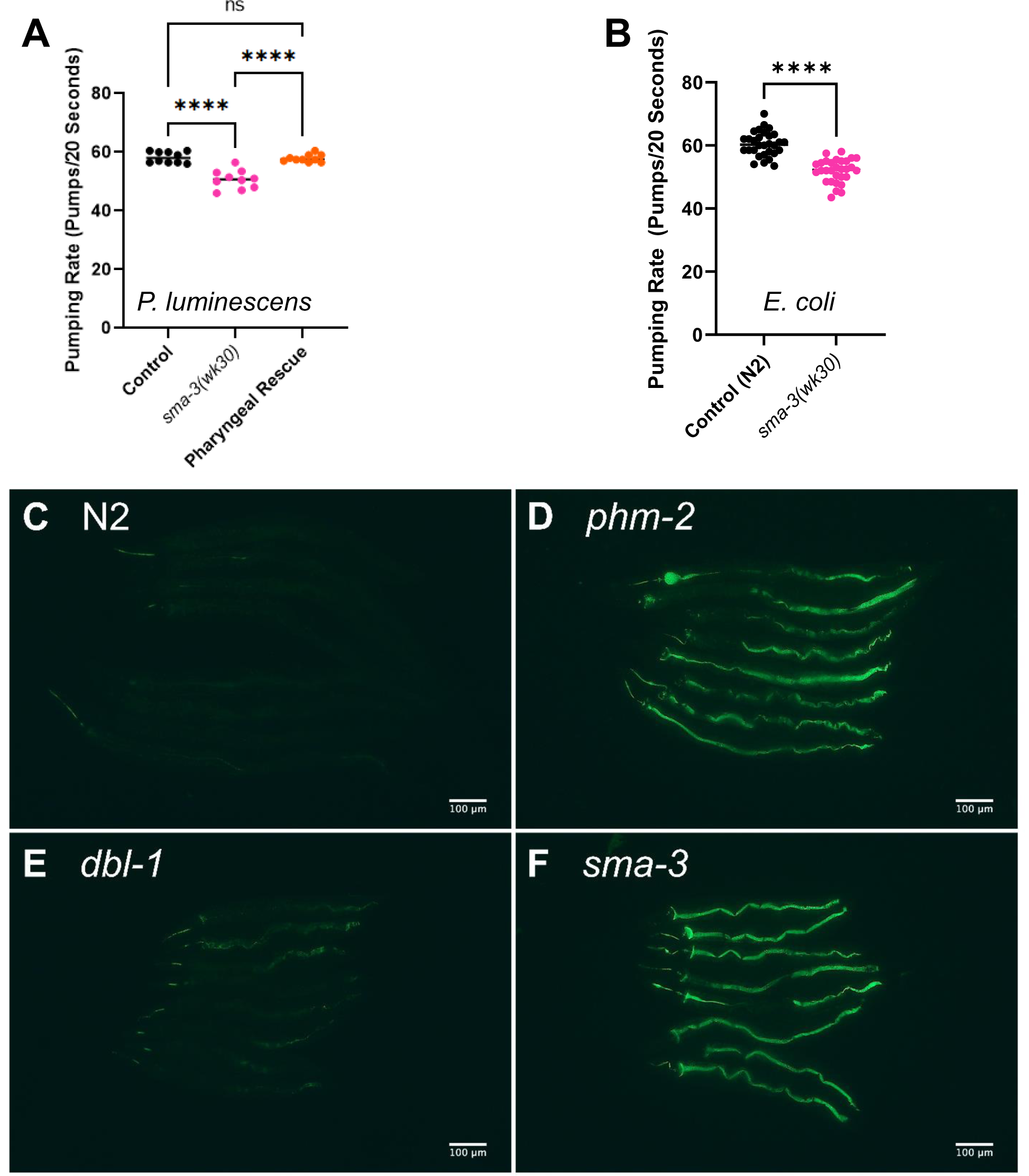
SMA-3 controls pharyngeal pumping rate cell autonomously. **A.** Pumping rate in pumps/20 Seconds of *sma-3(wk30)* mutants and pharyngeal *sma-3* rescue animals on *P. luminescens* bacteria. **B.** Pumping rate in pumps/20 Seconds of *sma-3(wk30)* mutants and on *E. coli* bacteria. 10 worms per strain were counted and experiments were repeated on two-three separate biological replicates. Results were consistent across all trials. Statistical analysis was performed using One way ANOVA with Šídák’s multiple comparison test, using GraphPad Prism 9. ns P > 0.05; **** P **≤** 0.0001. **C-F.** L4 animals of the designated genotypes were exposed to fluorescent OP50-GFP *E. coli* bacteria for 24 hours and 10 or more animals imaged as Day 1 adults. The experiment was repeated in triplicate. All scale bars show 100 µm.

To assess whether functionality of the pharynx contributes to survivability on pathogenic bacteria, we determined the survival patterns and pumping rates of two pharyngeal defect mutants. We used *eat-2* and *phm-2,* both involved in pharyngeal grinder function, for these experiments. Mutations of *eat-2* (Avery, 1993; Raizen et al., 1995; Lakowski and Hekimi, 1998; Avery and You, 2012) or of *phm-2* (Avery, 1993; Kumar et al., 2019; Scharf et al., 2021) result in reduced grinder function and a reduced rate of pharyngeal pumping. Following exposure of either strain to *P. luminescens* bacteria for 24 hours, there was a significant decrease in pharyngeal pumping rate (Fig 6AB), consistent with our expectations. Furthermore, both *phm-2* and *eat-2* mutants have a significantly reduced rate of survival compared to control animals when exposed to *P. luminescens* as well (Fig 6CD). Other studies have also demonstrated that both mutations result in sensitivity to bacterial infection, due to live bacteria reaching the intestine for colonization (Labrousse et al., 2000; Smith et al., 2002; Portal-Celhay et al., 2012; Kumar et al., 2019; Scharf et al., 2021). Taking all of our results together with these prior studies suggests that disrupting pharyngeal function contributes to susceptibility to infection.

**Figure 6.**
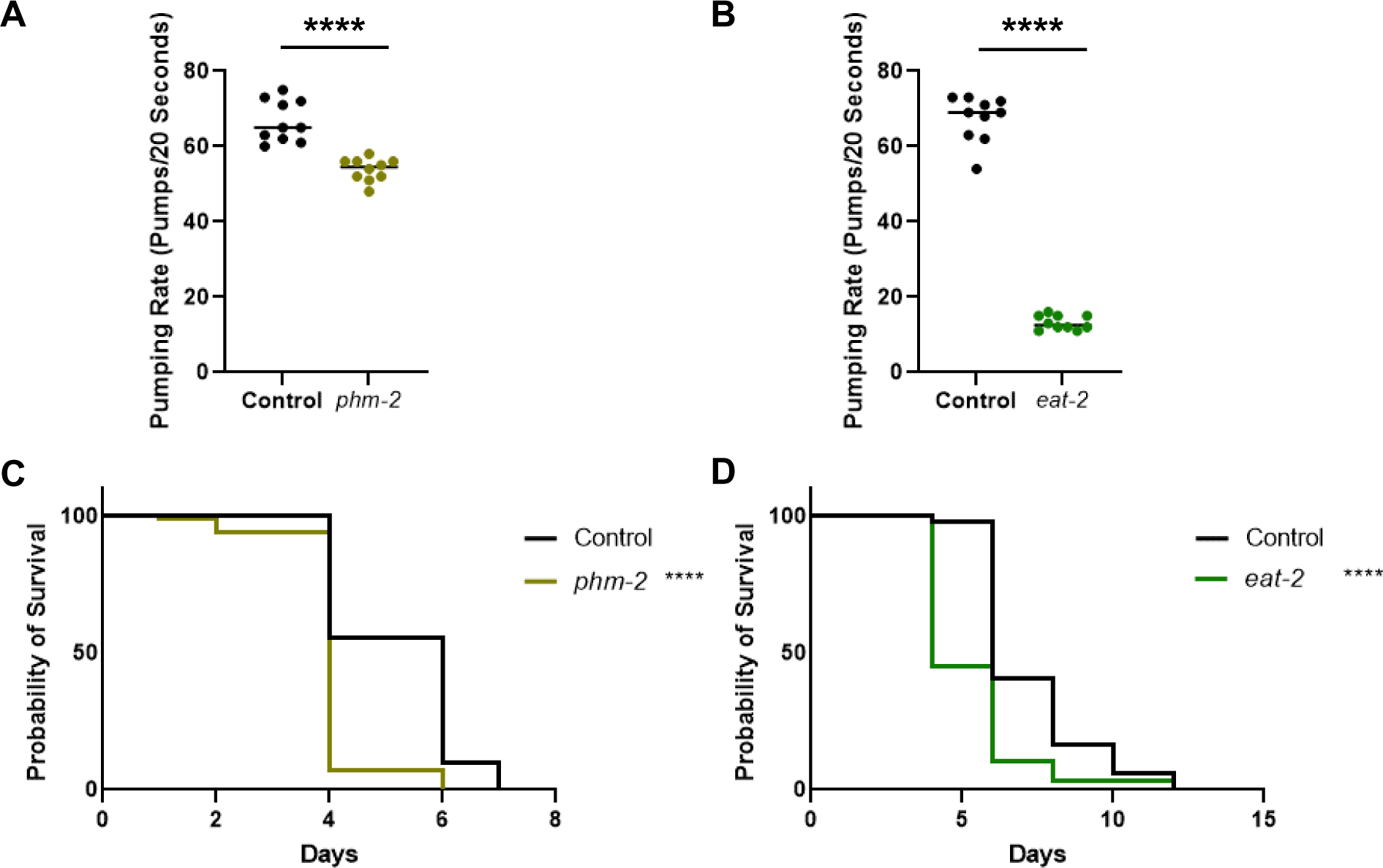
Mutants with pharyngeal defects have reduced survival on *P. luminescens* bacteria. **A-B**. Pharyngeal pumping rate in pumps/20 Seconds for pharyngeal defect mutants *phm-2(ad597)* and *eat-2(ad1116)* on *P. luminescens* bacteria. 10 worms per strain were counted and experiments were repeated on two separate biological replicates. Results were consistent across all trials. Statistical analysis was performed using One way ANOVA with Šídák’s multiple comparison test, using GraphPad Prism 9. **C-D.** Survival analysis of *phm-2(ad597)* and *eat-2(ad1116)* on *P. luminescens* bacteria. N values: Control (81), *phm-2* (99), Control (96), *eat-2* (58). Statistical analysis was done using Log-rank (Mantel-Cox) Test. **** P **≤** 0.0001.

## Discussion

The BMP-like ligand DBL-1 has previously been shown to play a significant role in the *C. elegans* immune response to bacterial pathogen (Mallo et al., 2002). Additionally, it has been demonstrated that the DBL-1 pathway effector, R-Smad SMA-3 is involved in a non-canonical signaling response to fungal infection, in which it acts without SMA-2 or the Co-Smad SMA-4 (Zugasti and Ewbank, 2009). A strength of the *C. elegans* system is the ability to ask questions about site of action in the context of the intact organism, shedding light on how systemic organismal responses may be generated. We had previously shown that *sma-3* expressed solely in the epidermis is sufficient to restore normal body size in adult animals (Wang et al., 2002). In this study, we employed survival analyses to determine where SMA-3 acts in the *C. elegans* immune response to bacterial pathogens. We demonstrate that *sma-3* expression in the intestinal site of infection does not have any measurable effect on survival against bacterial pathogen. However, expression of *sma-3*, and therefore pathway activity, in the pharynx, does restore survival towards that of control animals. Improved survival of pharynx-specific expressing strains as compared to *sma-3* mutants suggests the cell non-autonomous activity of the DBL-1 pathway on immunity.

We also determined whether the effect of DBL-1 signaling on the immune response to bacterial pathogen is related to regulation of immune response related genes. Here we demonstrate through qRT-PCR analysis that DBL-1 signaling regulates the expression of *abf-2* and *cnc-2* transcripts upon exposure to pathogenic *P. luminescens*. We also show that this effect is regulated through R-Smad SMA-3 expression in the pharynx. This result supports the hypothesis that DBL-1 signaling to regulate AMP expression is both cell autonomous and cell non-autonomous. We note that expression levels in some of the *sma-3* transgenic strains are higher than those seen in N2 control animals, which could be due to a higher level of expression of SMA-3 from the multi-copy transgenes.

The antibacterial factor related peptide ABF-2 has previously been shown to play a role in the response to bacterial pathogen (Kato et al., 2002). *abf-2* has a complex relationship with *abf-1:* the two genes are capable of being transcribed as a polycistronic operon transcript, but also as individual genes with different expression patterns. It will be interesting to determine whether SMA-3 acts at the level of transcription or post-transcriptionally to mediate regulation of these genes. In contrast, CNC-2 has been primarily correlated with fungal infection of the epidermis (Zugasti and Ewbank, 2009; Zehrback et al., 2017) and has not previously been associated with induction in response to *P. luminescens* infection. Previous investigations utilizing microarray and RNA-seq have only found decreases in *cnc-2* upon *P. luminescens* infection, although this difference may relate to the different methodologies used (Wong et al., 2007; Engelmann et al., 2011). In light of this, we also looked to see how *cnc-2* mutants survive on *P. luminescens* and found that *cnc-2* mutant animals had a significantly reduced survival compared to control animals. This increased susceptibility is consistent with our findings by qRT-PCR and the results demonstrating that DBL-1 signaling regulates *cnc-2* expression levels in response to bacterial pathogen exposure as part of the *C. elegans* immune response to *P. luminescens* infection.

Because the focus of action of SMA-3 in innate immunity is the pharynx, we considered whether DBL-1 signaling activity in the pharynx influences the physical function of the pharynx. The pharynx itself is an early site of exposure for *C. elegans*, as ingestion of bacteria brings pathogen through the pharynx before intact bacteria can be passed to the intestine. Typically, the pharyngeal grinder functions to break down these bacterial cells mechanically, inhibiting live bacteria from entering the intestine, preventing infection, and providing nutritional support. However, in instances of reduced pharyngeal pumping/grinder activity or exposure to more pathogenic bacteria, elevated levels of live bacteria in the intestine result in proliferation and infection (Kurz et al., 2003; Gravato-Nobre and Hodgkin, 2005).

We used pharyngeal pumping rate as a method for measuring the effect of *sma-3* on pharyngeal mechanical action (Raizen et al., 2012). We found that *sma-3* mutant animals have a reduced pharyngeal pumping rate as compared to control animals and that this reduction can be rescued with expression of *sma-3* in pharyngeal muscle. Similarly, pharyngeal function mutants *phm-2* and *eat-2* have reduced survival on *P. luminescens.* We demonstrated that *sma-3* mutants and *phm-2* mutants accumulate fluorescent *E. coli* in the intestine, demonstrating that reduced pharyngeal pumping can lead to diminished physical disruption of bacteria. In the case of pathogens, increased levels of live bacteria entering the intestine can cause pathological harm. In summary, we have identified the pharyngeal muscle as a critical responder to BMP signaling to generate a system organismal response to pathogenic bacteria. R-Smad SMA-3 in the pharynx is sufficient to rescue multiple defects, including reduced survival, loss of induction of AMPs, and altered pharyngeal pumping. Although most work on the *C. elegans* immune response to bacteria has focused on the intestine, the pharynx is an earlier point of contact for exposure to bacteria and may play a more important role than previously recognized.

## Materials and Methods

### Nematode Strains and Growth Conditions

*C. elegans* were maintained on *E. coli* (DA837) at 20°C on EZ worm plates containing streptomycin (550 mg Tris-Cl, 240 mg Tris-OH, 3.1 g Bactopeptone, 8 mg cholesterol, 2.0 g NaCl, 200 mg streptomycin sulfate, and 20 g agar per liter). The N2 strain was used as a control unless otherwise specified. In tissue-specific *sma-3* experiments, the strains used were: CS152 *qcIs6[GFP::sma-3* + *rol-6(su1006)]*, CS640 *sma-3(wk30); qcIs53[myo-2p::*GFP*::sma-3 + rol-6(su1006)],* CS619 *sma-3(wk30); qcIs59[vha-6p::*GFP*::sma-3 + rol-6(su1006)]*, CS215 *sma-3(wk30); him-5(e1490); qcEx55[vha-7p::GFP::sma-3* + *rol-6(su1006)],* CS635 *sma-3(wk30); qcIs62[rol-6(su1006)]*. Remaining strains used in this study were*: sma-3(wk30), cnc-2(ok3226)*, *sma-6(wk7)*, *sma2(e502)*, *dbl-1(wk70), phm-2(ad597), eat-2(ad1116).* All mutations employed are strong loss-of-function or null alleles. In particular, the *sma-3* allele *wk30* contains a premature termination codon.

### Bacteria

Control bacteria used in all experiments was *E. coli* strain DA837, cultured at 37°C. *S. marcescens* strain Db11 cultured at 37°C and *P. luminescens* (ATCC #29999) cultured at 30°C were used for pathogenic bacteria exposure. *Serratia* (Db11) was seeded on EZ worm plates containing streptomycin and grown at 37°C. *P. luminescens* was seeded on EZ worm plates with no antibiotic and grown at 30°C.

### Survival Analysis

Survival plates were prepared at least one day prior to plating worms. Each plate was seeded with 500 µl of pathogenic bacteria. FuDR (5-Fluoro-2’-deoxyuridine) was used to prevent reproduction, at a concentration of 50 µM per plate. This treatment was done to reduce the incidence of matricide by internal hatching of embryos during survival analysis. All survival experiments were carried out at 20 °C. All survival analyses were repeated. All graphs were made using GraphPad Prism software and statistical analysis performed using Logrank/Mantel Cox test.

### qRT-PCR Analysis

Worms were synchronized using overnight egg lay followed by four-hour synchronization. When animals reached L4 stage, they were washed and moved to experimental plates for 24-hour exposure to *P. luminescens*. Animals were collected and washed after 24-hour exposure and RNA was extracted using previously published protocol (Yin et al., 2015) followed by Qiagen RNeasy miniprep kit (Cat. No. 74104). cDNA was generated using Invitrogen SuperScript IV VILO Master Mix (Cat. No.11756050). qRT-PCR analysis was done using Applied Biosystems *Power* SYBR Green PCR Master Mix (Cat. No. 4367659). Delta delta Ct analysis was done using Applied Biosystems and StepOne software. All qRT-PCR analysis was repeated on separate biological replicates. All graphs were made using GraphPad Prism software and statistical analysis was performed using One-way ANOVA with Multiple Comparison Test, as calculated using the GraphPad software.

### Pumping Rate Experiments

Pumping rate experiments were done with animals exposed at L4 stage for 24 hours on *P. luminescens* bacteria. Pharyngeal pumps were counted over 20 second timespan (Raizen et al., 2012). All pumping experiments were repeated. Graph made using GraphPad Prism software and statistical analysis performed using One-way ANOVA with Multiple Comparisons Test, as calculated using the GraphPad software.

### OP50-GFP Assay

Antibiotic-free plates were each seeded with 500 µL of OP50-GFP and incubated overnight at 37℃. 20 L4 animals were picked onto each OP50-GFP plate. After 24 hours, worms were prepared for imaging. Worms were picked into a droplet of M9 on a regular OP50 plate and allowed to crawl out of the droplet. This removed any surface fluorescent bacteria left on the animal, which may interfere with imaging. Worms were mounted on 2% agarose pads containing a 5 µL drop of 2.5 mM levamisole for immobilization. A small hair was used to align the animals. Images were taken on a Zeiss Axioskop with Gryphax software and a 10X objective. Experiment was repeated in triplicate.

### Imaging GFP Reporter Strains

L4 animals were mounted on 2% agarose pads containing a 5 µL drop of 2.5 mM levamisole for immobilization. Images were taken on a Zeiss ApoTome 3 with Zen Pro software and a 20X objective.

## ACKNOWLEDGEMENTS

This work was funded by NIH awards R15GM112147 and R21AG075315, and by a PSC-CUNY grant to CSD. Some strains were provided by the Caenorhabditis Genetics Center, which is supported by the National Institute of Health - Office of Research Infrastructure Programs (P40 OD010440). The *cnc-2(ok3226)* strain was generated by the *C. elegans* Gene Knockout Project at the Oklahoma Medical Research Foundation, part of the International *C. elegans* Gene Knockout Consortium. Genetic data from WormBase (Davis et al., 2022) were critical to the execution of this project. This work was carried out in partial fulfillment of the requirements for the Ph.D. degree from the Graduate Center of City University of New York (EJC and KKY).

## Supporting Information

**S1Fig.**
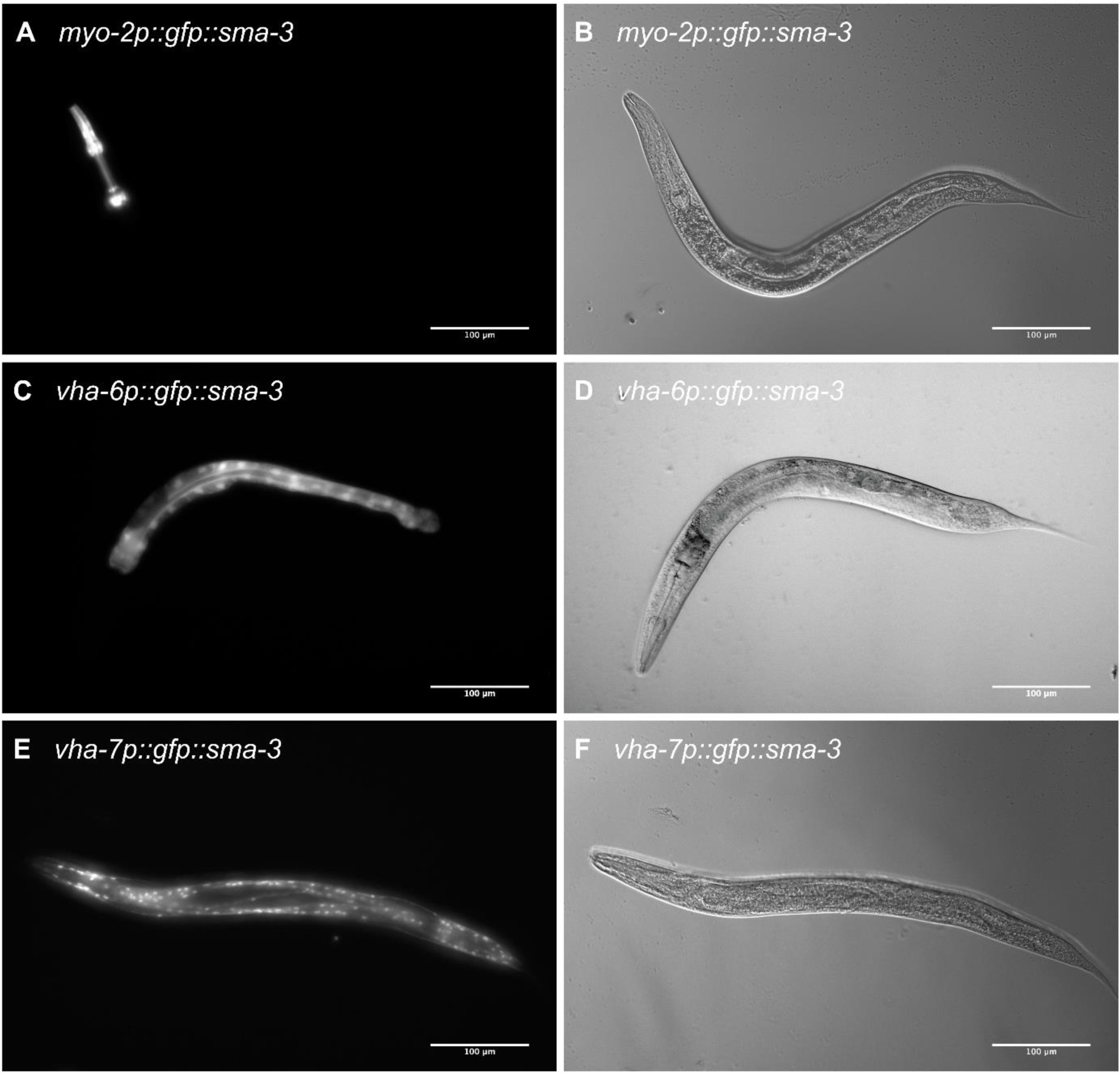
Representative images of *sma-3* transgenic strains. Fluorescence (**A**) and DIC (**B**) images of *myo-2p::GFP::SMA-3* demonstrating specific GFP::SMA-3 expression in pharyngeal muscle. Fluorescence (**C**) and DIC (**D**) images of *vha-6p::GFP::SMA-3* demonstrating specific GFP::SMA-3 expression in intestine. Fluorescence (**E**) and DIC (**F**) images of *vha-7p::GFP::SMA-3* demonstrating specific GFP::SMA-3 expression in epidermis. Scale bars show 100 µm.

**S2Fig.**
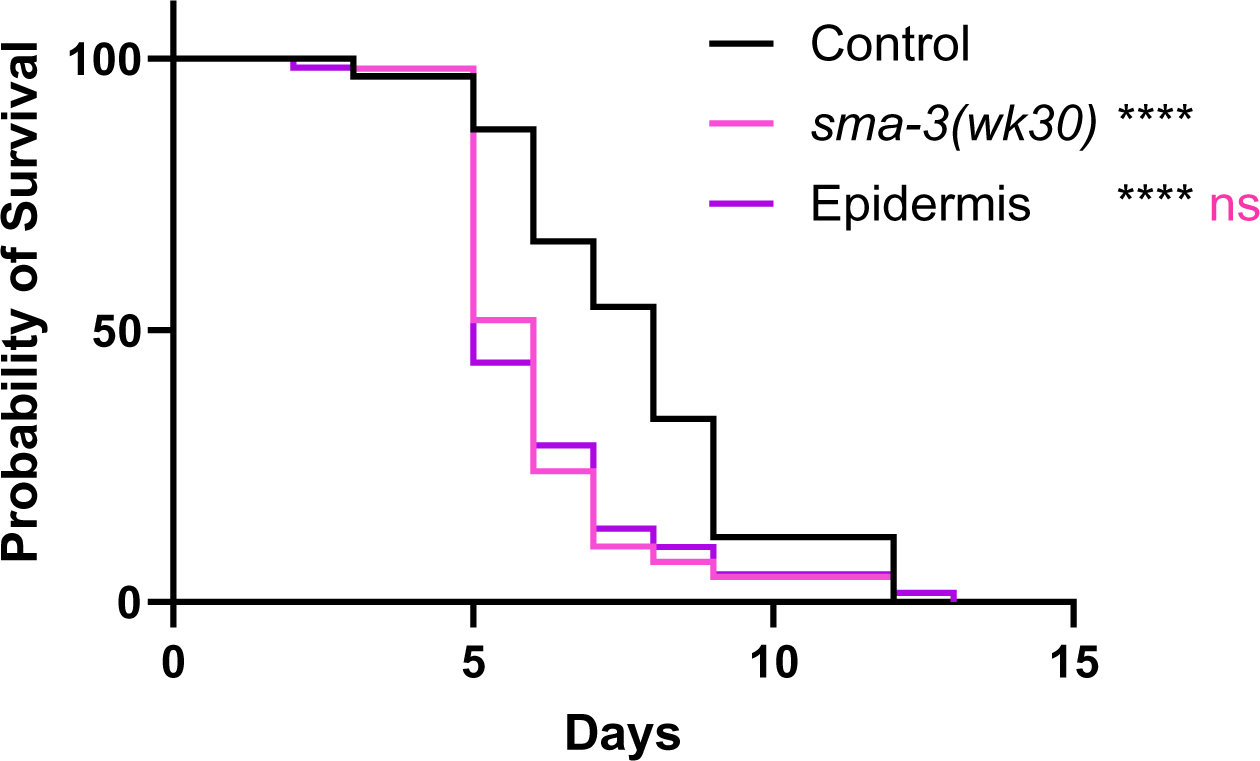
Survival of strain with epidermal-specific *sma-3* expression. **A.** CS215 carrying an extrachromosomal array driving expression of *sma-3* in the epidermis via the *vha-7* promoter (Supporting Figure S1) was analyzed. n values on *P. luminescens.* n values: Control (92), *sma-3* (108), Epidermis (59). Survival analyses were repeated and results were consistent across trials. Statistical analysis was done using Log-rank (Mantel-Cox) Test. ns P > 0.05; **** P < 0.0001. Black asterisks show significance relative to control; magenta shows significance relative to *sma-3* mutant.

**S3Fig.**
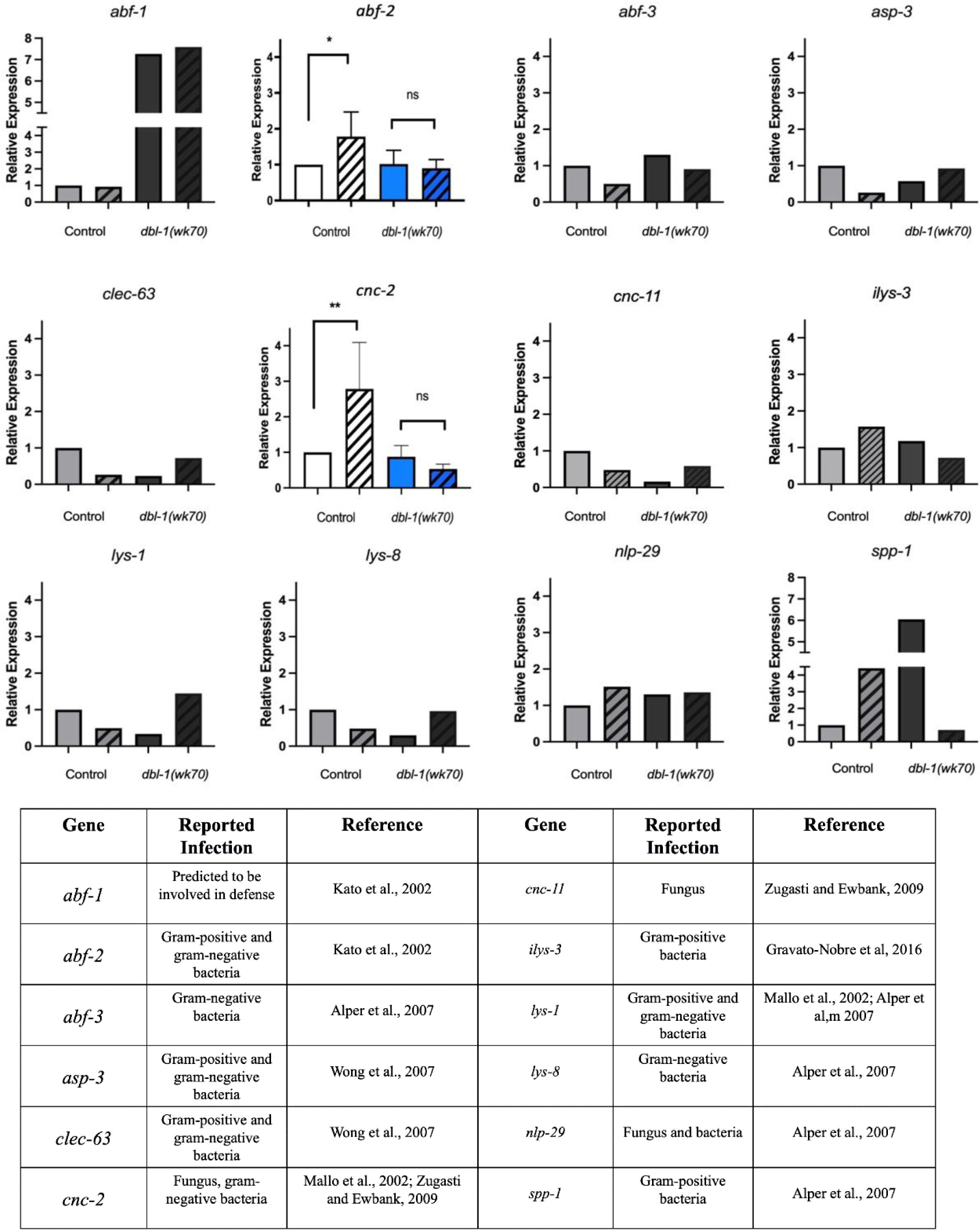
qRT-PCR analysis panel of 12 immune response genes in *dbl-1(wk70)* mutants. qRT-PCR analysis of 12 immune response genes in control and *dbl-1(wk70)* mutant animals, comparing 24-hour bacterial pathogen exposure (diagonally hatched bars) to control conditions. *P. luminescens* was used as the bacterial pathogen. Data for *abf-2* and *cnc-2* represents repeated analyses for two biological replicates. The remaining genes were analyzed for one biological sample. Graph was made using GraphPad Prism. Statistical analysis for *abf-2* and *cnc-2* was done using One-way ANOVA with Šídák’s multiple comparison test, using GraphPad Prism 9. ns P > 0.05; * P ≤ 0.05; ** P ≤ 0.01.

